# Nanostructure of the desmosomal plaque

**DOI:** 10.1101/2022.04.18.488620

**Authors:** Irina Iachina, Helle Gam-Hadberg, Jonathan R. Brewer

## Abstract

Desmosomes are considered one of the most important intercellular junctions with respect to mechanical strength. Therefore, their spatial distribution and structure is of interest with respect to understanding both healthy and diseased tissue. Previous studies have imaged desmosomes in tissue slices using transmission electron microscopy, or low-resolution confocal images, but both these techniques lack the ability to resolve the 3-dimensional structure of the desmosomes. In this work it was possible to determine the 3D-nanostructure of single desmosomal complexes in both mouse and human epidermis, by 3D stimulated emission depletion (STED) microscopy. STED images of desmoplakin and the desmosomal cadherin, desmoglein revealed that desmosomes form ring-like structures, distributed over the cell surface, with diameters of around 1 μm. STED images of the tight junction plaque protein ZO1 also displayed ring formations, suggesting a common structure for intercellular junctions. Measurements of the desmosomal plaque protein, desmoplakin showed an increased intercellular plaque distance during the stratum basale (0.23±0.027µm) to stratum spinosum (0.28±0.039 µm) transition.

## Introduction

Desmosomes are a class of intercellular junction plaques, together with the other intercellular junction plaques such as gap junctions and tight junctions [1]. Desmosomes are specialized cell-cell adhesion complexes that provide tissue with resilience and strength, by linking the keratin intermediate filament of adjacent cells (Figure 1).

**Figure 1:**
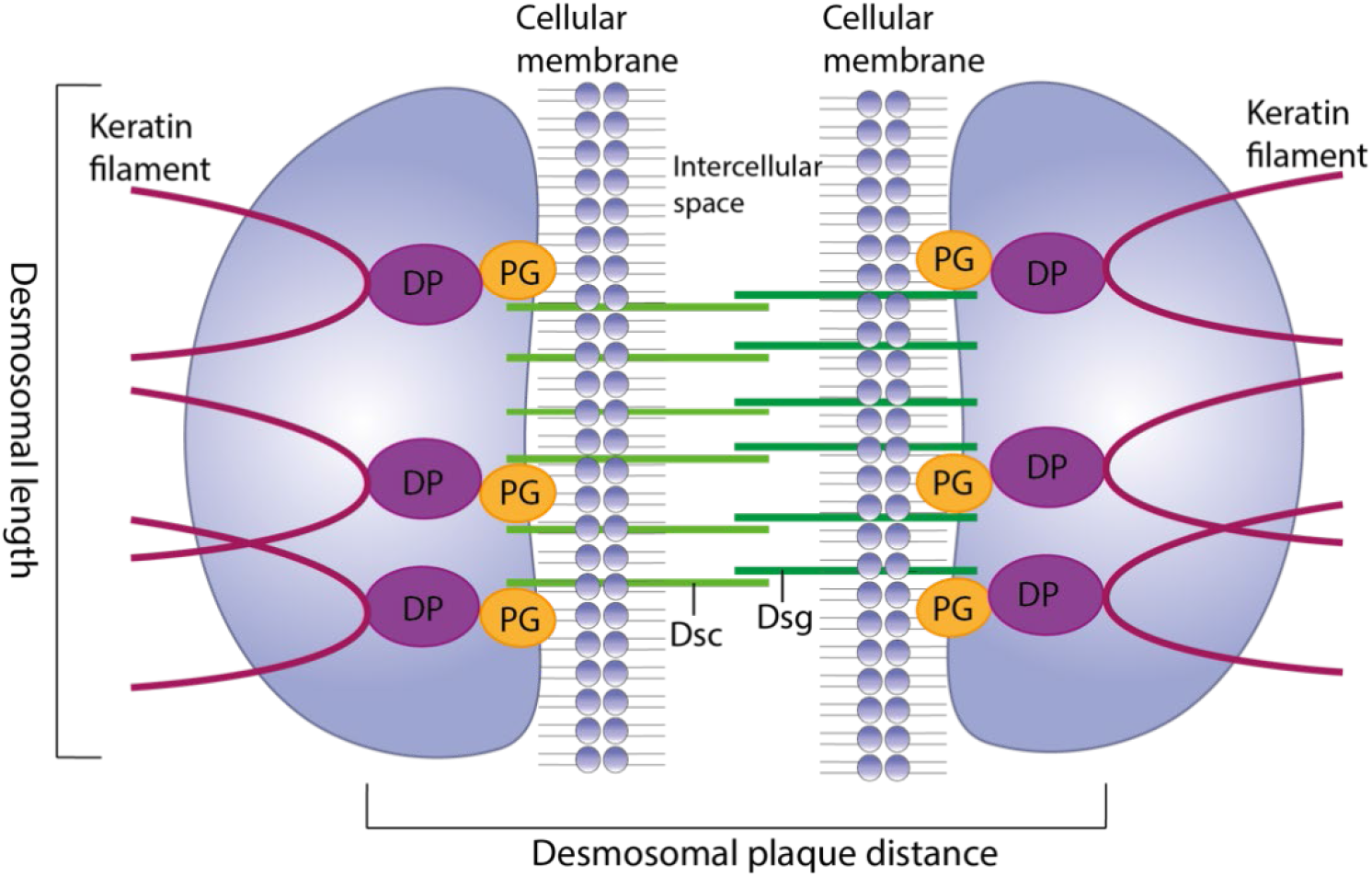
Schematic representation of the desmosomal plaque. In the intracellular space the plaque consists of desmoplakin (DP) and plakoglobins (PG), and the former binds to keratin filaments. The desmosomal cadherins desmocollin (Dsc) and desmoglein (Dsg) anchor the two neighboring cells. Shown is the desmosomal plaque distance as the distance between the C-terminals in DP in two neighboring plaques, as well as the desmosomal length.

Thus, these complexes are often seen in tissues that undergo regular mechanical stress, such as epithelia, cardiac muscle, bladder, and intestines [2-4]. Disruption of the desmosomes can lead to disturbance of the epidermal homeostasis, a reduction in epidermal barrier function, blistering disease and overall skin fragility due to decreased intercellular attachment [5-7]. Their spatial distribution and structure are therefore of great interest both for understanding healthy and diseased tissue.

Desmosomes were first discovered on the surface of epidermal cells by Schrön in 1863[8] who interpreted them as pores in the cellular membrane but were later acknowledged as an intercellular junction, bridging the intercellular space, by Bizzozero in 1870 [9]. In 1958 Odland[10] reported the desmosomes to be disc-shaped, and there has since been a general agreement on the desmosomal structure. However, the reported diameter and width of desmosomes has varied greatly and early confusion identifying the various intercellular junctions has made the understanding of the desmosomal structure even more unclear [10-13].

Typically, desmosomes have been imaged in tissue slices using Transmission Electron Microscopy (TEM) [14-16], or low-resolution confocal images [17]. However, both these techniques lack the ability to resolve the 3-dimensional structure of the desmosomes.

Super resolution optical microscopy, such as Stochastic Optical Reconstruction Microscopy (STORM) [18] and stimulated emission depletion (STED) microscopy, have been used to provide ultra-structural images of intact tissue, where a resolution down to 40-50 nm was achieved compared to a resolution of 200 nm using regular fluorescence microscopy [19-22]. Recently, STORM has been used to resolve the structures of the desmosomes in tissue reveling them as parallel rows of plaques on each side of the intercellular gap [13, 23, 24]. Interestingly, from literature it is known that gap junctions form round pores in the cellular membrane of neighboring cells, to establish a route of communication between neighboring cells [25]. However, the 3-dimensional structure of the desmosomes and tight junction plaques are as yet not fully elucidated. Therefore, in this study STED and confocal microscopy were used to investigate and visualize the 3D nanostructure of desmosomes. This was done by imaging the desmosomal plaque made up by the desmosomal plaque protein, desmoplakin (DP), as well as the desmosomal cadherin desmoglein (Dsg), and tight junction plaque protein ZO1 in human and mouse skin samples.

## Results and Discussion

For STED microscopy the protein desmoplakin (DP) was labeled in both human and mouse epidermis (Figure 2).

**Figure 2:**
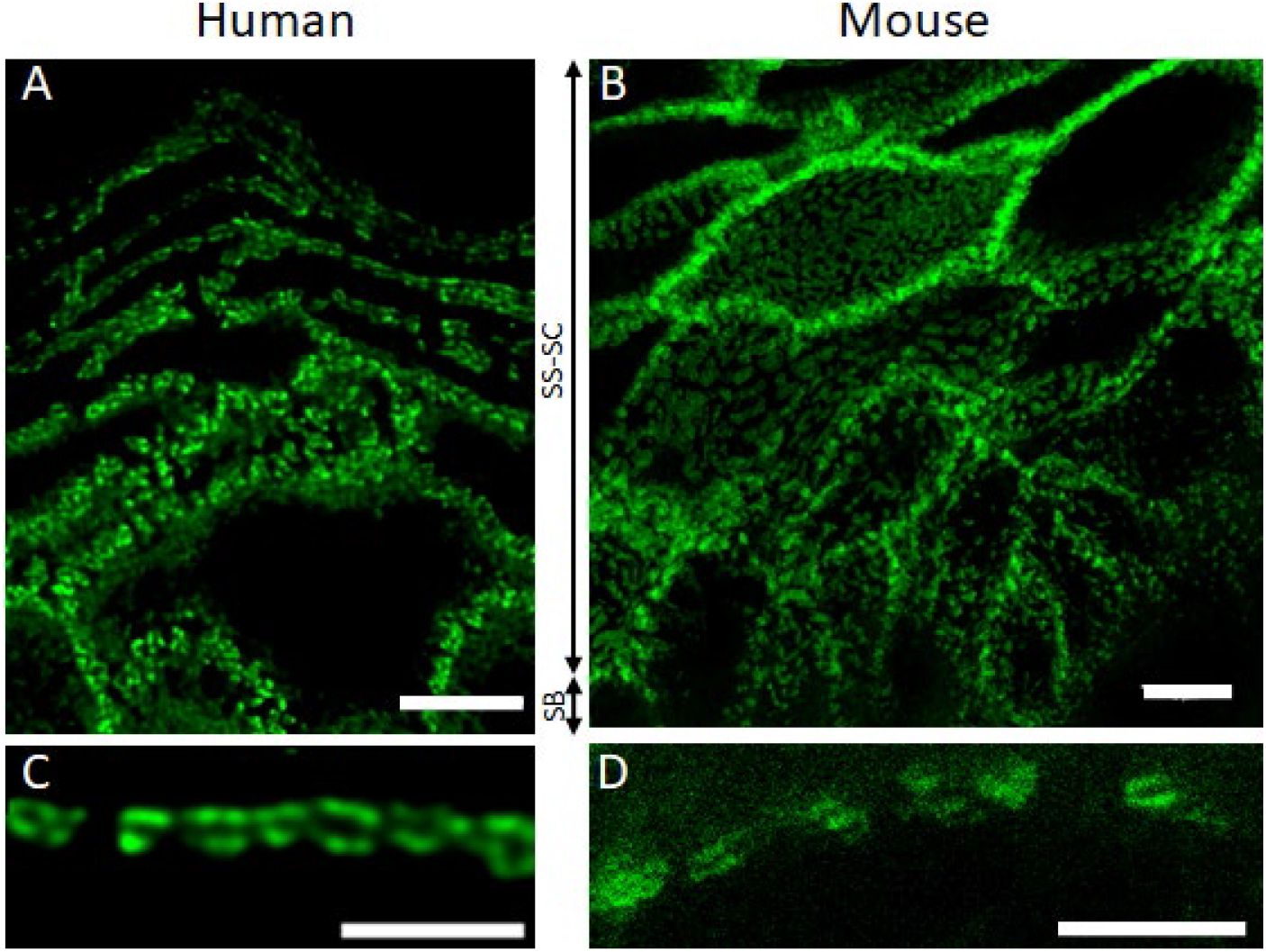
Desmosomal complexes outline the individual keratinocytes. A) and B) STED images of desmoplakin in human and mouse epidermis, respectively. The desmosomal plaques are located along the cellular membrane in a discontinuous pattern, outlining the individual keratinocytes in all the viable layers (SS-SC). The number of complexes decrease when entering the basal layers (SB). Scalebar is 5 μm. C) (Scalebar is 2 μm) and D) (Scalebar is 1 μm) STED images of desmoplakin in individual cells in human and mouse epidermis, respectively. One can clearly see the individual plaques laying opposite each other representing the two halves of a desmosome. The dark space between two plaques represents the intercellular proteins and the cellular membranes of the adjacent keratinocytes. Scalebar is 2 μm.

With STED microscopy (Figure 2C and D) a resolution of 60 nm was obtained, enabling the imaging of sub-resolution protein organizations and the 3-dimensional structure of the individual desmosomal components.

The desmosomes were found to be located along the cellular membrane outlining individual keratinocytes. As expected, the protein is expressed in all epidermal layers, with a slightly lower expression in the basal layer. The mirror symmetry of the desmosomal plaques (Figure 2C and D) makes it easy to identify the individual desmosomes along the cellular membrane. Here the two opposite plaques show intensive labeling separated by a dark gap, representing the intercellular space where the desmosomal cadherins, desmocollin (Dsc) and desmoglein (Dsg), as well as the cellular membranes of\ the neighboring cells reside. Even though the desmosomes are distributed along the cellular membrane of the entire cell, the individual desmosomes do not appear to have any physical contact.

Distances between the plaques were measured in both mouse and human epidermis, to determine desmosomal plaque distance, as illustrated in Figure 1. The plaque distances measured varied according to epidermal location, thus desmosomes connecting basal (SB) keratinocytes had a tighter configuration compared to desmosomes located in the suprabasal (SP) layers (Figure 3A). In desmosomes connecting basal cells the mean plaque distances were measured to be 0.23 µm (±0.027 μm SD, n=26) and 0.22 μm (±0.034 μm SD, n=57), for mouse and human epidermis respectively. In the suprabasal layers the mean distance measured was significantly wider (P-values can be seen in Table 1), as it was measured to be 0.28 µm (±0.034 μm SD_mice_, n=26, ±0.039 μm SD_human_, n=57) in both mouse and human epidermis.

**Table 1:** Shows the plaque distances (as shown in Figure 1), standard deviation, P-value and n samples measured using STED microscopy in basal and suprabasal cells in human and mouse epidermis. The P-values indicate that there is a statically significant difference between the plaque distances measured in the SB and the SP, both for mouse and human.

| Sample | Plaque distance<br>[$\mu\text{m}$] | Standard deviation [$\mu\text{m}$] | P-value | n |
| --- | --- | --- | --- | --- |
| Basal cells, human | 0.22 | $\pm 0.034$ | | 26 |
| Suprabasal cells, human | 0.28 | $\pm 0.039$ | 7.4E-7 | 26 |
| Basal cells, mouse | 0.23 | $\pm 0.027$ | | 57 |
| Suprabasal cells, mouse | 0.28 | $\pm 0.034$ | 2.5E-13 | 56 |

**Figure 3:**
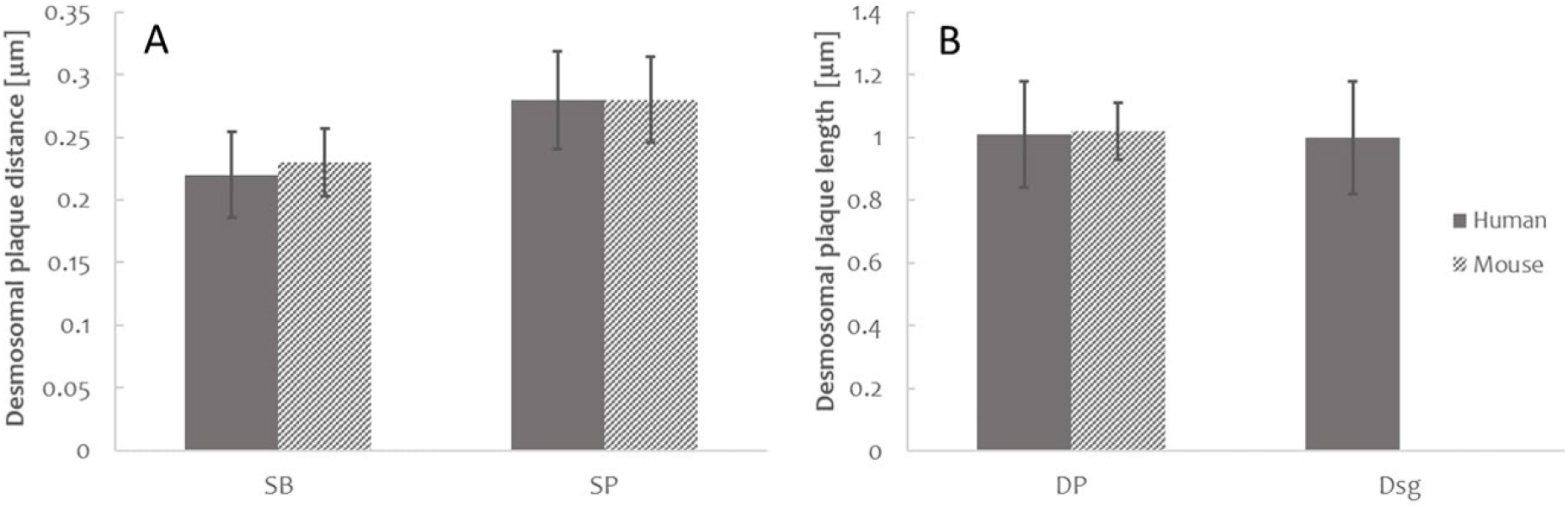
STED enabled exact measurements of the desmosomal dimensions. A) Plaque distances [μm] measured (as shown in Figure 1) in basal (SB) and suprabasal (SP) layers in human and mouse epidermis. The measurements revealed a significantly shorter plaque distance (PD) in basal keratinocytes (PD_human_=0.22 μm ± 0.034, PD_mouse_ = 0.23 μm ± 0.027) compared to suprabasal keratinocytes (PD_human_=0.28 μm ± 0.039, PD_mouse_ = 0.28 μm ± 0.034, in both human and mice. P-values can be seen in Table 1. B) Desmosomal length (diameter) [μm] measured (as shown in Figure 1) in STED images when desmoplakin, DP (1 μm ± 0.17) or desmoglein, Dsg (1 μm ± 0.18) was labeled. No significant difference was found in the diameter of the desmosome when measuring the intercellular and extracellular part of the complex (P=0.7, n_DP_=37, n_Dsg_=7)

The wider configuration in the suprabasal layers could be associated with the maturation of the desmosomes during which the desmosomes switch from a Ca^2+^-dependent state to a hyper-adhesive Ca^2+^-independent state. This was also hypothesized by Stahley et al. [13], who also observed a tighter configuration in desmosomes connecting basal keratinocytes compared to desmosomes found in suprabasal keratinocytes. During wound healing the hyper-adhesive state is reversed to more plastic Ca^2+^-dependent state to facilitate cell migration [13, 26]. Hence the wider structure would represent a more rigid structure optimized for distributing forces along the keratin intermediate filaments, when exposed to mechanical stress.

Alternatively, the change in expression of the desmosomal cadherins could be responsible for the change in desmosome width. All seven desmosomal cadherins are expressed in the epidermis, but as the keratinocytes undergo terminal differentiation, their expression level changes. Dsg2 and 3 as well as Dsc2 and 3 are mostly observed in the lower epidermal layers, while Dsg1 shows increased expression up through the epidermal layers and Dsc1 has the highest expression in the granular layer [27, 28]. Since the desmosomal cadherins are differentially expressed, desmosomes are biochemically, and therefore possibly, biologically different throughout the layers[6]. But since the change in desmosomal cadherin expression is gradual and the change in desmosomal configuration is prompt when entering the spinous layer, the change from a Ca^2+^-dependent state to a Ca^2+^-independent hyper-adhesive state seems the most likely explanation.

As mentioned earlier, the reported length of the desmosomes has also varied greatly. Here, the mean length of the desmosomal plaque measured parallel to the cellular membrane as shown in Figure 1, was approximately 1 µm in both human and mouse epidermis (Figure 3B) and nearly twice the length previously reported by Odland [10] and Selby [13, 29]. To make sure the diameter of the plaque represented the diameter of desmosomes in general, the diameter of the intercellular domain, where desmoglein is present was labeled and measured. The diameter of the intercellular domain did not vary significantly from the previous measurements (Figure 3B). The differences in the dimensions reported could partly be related to the sample preparation. An observation supported by Farquhar and Palade [12] as well as Bates et al [30], the former of which reported that choice of fixative affects the thickness of the cellular membrane, and the latter reported the use of primary and secondary antibodies affect the resolution and will increase the apparent size of structures in the tissue. This of course does not entirely explain the discrepancies and one could imagine that the structure of the desmosomes could be tissue specific. This in turn raises the question whether the size of the desmosomes is directly related to strength or rigidness, so tissues that undergo similar levels of mechanical stress have similar sized desmosomes.

To visualize the 3-dimensional structure of the desmosomes 3D STED z-stacks of the samples were recorded. STED images taken at the bottom of cells showing the plaques face on clearly showed ring shaped structures of the desmoplakin (Figure 4A). Further support for this ring structure comes from xz 3D STED images of the sides of cells which also showed clear ring structures (Figure 4B). Using STED microscopy, it was also possible to image single plaques, sliced through the side, located at the lateral part of the keratinocyte (Figure 4C and D). These images are typical of how desmosomes are depicted in optical microscopy in the literature, showing train track like structures of the plaques [13, 23, 24]. The STED images of the plaques located at the lateral part of the keratinocytes showed two distinct labeling patterns. The first is a continuous labeling pattern with a uniform intensity along the length of the entire plaque (Figure 4C). This pattern is consistent with the disc-shaped plaques reported by Odland (1958). The second pattern shows a non-continuous labeling pattern (figure 4D), with intensive labeling at the end and with no or faint labeling in the middle. These patterns do not reflect a disc-shaped structure of the desmosomal plaque. When dissecting a bowl- and ring structure (figure 4 E and F) only the ring-like structure can create the labeling pattern seen in figure 4D, with intense labeling at the ends and with no or faint labeling in the middle. This conformation corresponds to the ring being cut through the middle with only the ends of the ring giving signal to the image. Therefore, the structures seen in Figure 4C and D also indicate a ring like structure for the desmoplakin. For similar images in mouse see Figure S1.

**Figure 4:**
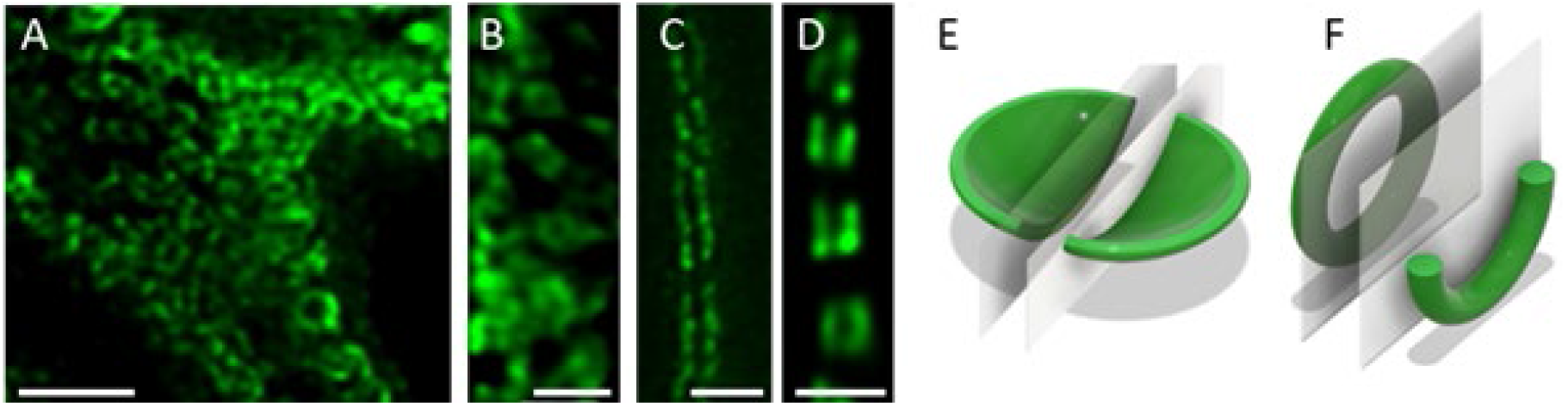
3D structure of desmosomal plaque. A-D) 3D STED images of keratinocytes with desmoplakin labeling in sliced human epidermis A) xy-plane image at the bottom of a keratinocyte where clear ring structures are seen. Scalebar is 2µm. B) xz-plane images of the side of a keratinocyte showing the desmosomal nanostructure resembles a ring. Scale bar 1 µm. C and D) B) xy-plane image of desmoplakin seen sliced through the side of the plaque, located at the lateral part of the keratinocyte. The labeling demonstrates the two distinct labeling patterns. In one the labeling continues along the entire length of the plaque (C), while a more discontinuous pattern with either no or very faint labeling in the middle but intense labeling at the ends is shown in (D). Scalebars are 1 μm. E and F) Illustrations of a dissected bowl (E) and ring (F) located at the cytoplasmic part of the cellular membrane. Observed from the middle only the ring-like structure can form the labeling pattern shown in D.

To exclude the applied antibody as a cause for this labeling pattern an alternative antibody targeting another epitope in desmoplakin was also used (data not shown). This labeling showed the same variation in labeling pattern, which excludes the antibodies as the cause of the ring shapes and the two distinct labeling patterns observed. The hypothesis of a ring structure was further strengthened when ring-like structures were observed when desmoglein was imaged (Figure 5A).

**Figure 5:**
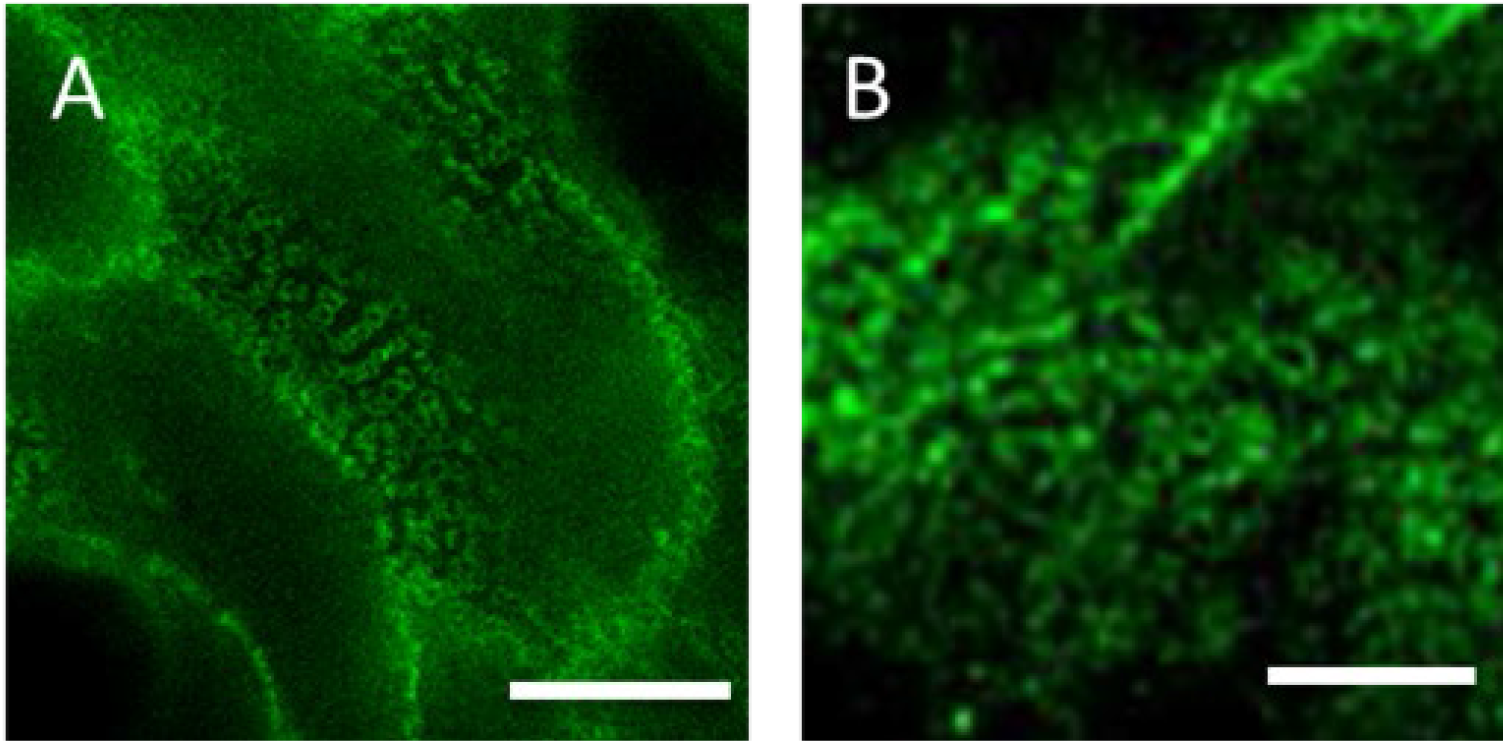
Possible uniform nanostructure for intercellular junctions. A) STED image of desmoglein in human epidermis. Image shows clear ring structure along the cellular membrane on the extracellular surface of the keratinocytes. Scalebar is 2 μm. B) STED image of zonula occludens 1 (ZO1) in mouse epidermis. Image reveals clear ring structures along the cellular membrane of the keratinocytes. Scalebar is 1 μm. Both observations strengthen the theory of a uniform structure for intercellular junctions.

Both the observed labeling pattern and the ring structure could explain the varied desmosomal sizes reported in literature. The absence of labeling in the middle could be interpreted as two individual desmosomes, resulting in smaller diameters.

Besides the desmosomes, there are several other intercellular junctions present in the epidermis. From the literature we know that gap junctions form round pores in the cellular membrane of neighboring cells, to establish a route of communication between neighboring cells [25]. Combined with our observations in the desmosomes, we hypothesize a common ring-like structure for intercellular junctions. To test the hypothesis, images of the tight junction plaque protein, zonula occludens 1 (ZO1), were taken. As one can see in Figure 5B, ZO1, just like desmoplakin, forms rings along the inner leaflet of the cellular membrane. This supports the suggestion of a common nanostructure for intercellular junctions but must of course be tested further in cadheren junctions as well as hemidesmosomes.

In summary, using STED microscopy it was possible to determine that desmosomes form a tighter structure in the basal layer compared to the suprabasal layers in both human and mouse epidermis. Whether there is a functional explanation to this conformational change or whether the change in tightness simply reflects the terminal differentiation program, needs further investigation. The increased resolution of 3D STED microscopy also revealed a ring-shaped nanostructure for the desmoplakin and desmoglein in the desmosomes, which are seen to coat the cells on all sides. STED images of the tight junction plaque protein ZO1 displayed similar ring formations, suggesting a common structure for the intercellular junctions such as desmosomes, tight junction as well as the gap junctions, however the nanostructure of the remaining junctions would need further investigation.

## Material and Methods

### Antibodies

Antibodies specific for desmoglein (ab16434) and ZO-1 (ab190085) were purchased from Abcam®, Cambridge, United Kingdom. Antibodies specific for desmoplakin I+II (ABIN452374) were purchased from antibodies-online.com, Aachen, Germany. Secondary antibodies coupled with STAR488 (rabbit: 2-0022-051-2; mouse: 2-0032-051-9) or STAR440SX (mouse: 2-0022-051-2; rabbit: 2-0032-051-9) and STAR-RED (Goat Anti Mouse, STRED-1001-500UG) were purchased from Abberior, Göttingen, Germany. Alexa Fluor 488 (A11055) was acquired from InvitrogenTM, Fisher Scientific, Slangerup, Denmark.

### Skin

Human skin samples were obtained from abdominoplasty and provided by the Department of Plastic Surgery at Odense University Hospital. where non-colored ethanol iodide was used for disinfection. Samples were stored at 4°C until processed, maximum 2h after surgery. The skin was cut into 0.5-1 thick 1 cm^2^ sections, washed in tap water before patted dry. Immediately afterwards the skin sections were coated in O.C.T compound (VWR, Søborg, Denmark) and frozen at -120°C in 2-methylbutan cooled with liquid nitrogen and stored at -80°C until use. According to Danish regulations human tissue remnants from surgery are regarded as waste products and hence do not require written informed consent from the patient. Any work involving human samples was performed in accordance with the Declaration of Helsinki Principles (2008).

The WT mice used in this study were provided by Thomas Magin and his group in Leipzig. Mice fetuses were removed by caesarean section on embryonic day 18.5. The mice were euthanized and frozen at - 120°C in 2-methylbutan cooled by liquid nitrogen. The samples where then send to us on dry ice and stored at -80°C until use.

### Labeling

Histological sections of 15 µm thickness were cut on the Cryotome FSE cryostat (Thermo Scientific, USA) and transferred to coverslips pre-coated with poly-L-lysin solution (1%) in H_2_O (Sigma-Aldrich, Copenhagen, Denmark). The tissue sections where fixed in -20 °C methanol for ten minutes followed by a blocking cycle (5×3 min) using phosphate buffered saline containing 1% bovine serum albumin (Sigma-Aldrich, Copenhagen, Denmark). The samples where then incubated overnight at 4 °C with the primary antibodies (dilution factor: Desmoplakin I+II 1:200, Desmoglein 1 1:500, ZO1 1:100). After incubation the sections were subjected to another rinse cycle (5×3 min) with phosphate buffered saline containing 1% bovine serum albumin and subsequently incubated overnight at 4 °C with the secondary antibodies. All samples were mounted on SuperFrost Plus microscope slides (Thermo Scientific, USA) using ProLong Diamond Antifade Mountant (Thermo Fisher, Hvidovre, Denmark)

### Imaging

Confocal and STED images of samples with STAR440SX, STAR488 and Alexa488 were obtained using a Leica TCS SP8 (Manheim, Germany) inverted stage microscope fitted with a Leica STED module (592 nm continuous wave depletion laser). Emission was recorded using a gated hybrid detector (0.3 ns.). The software used for controlling the setup of the microscope was Leica Application Suite X (LAS X). 3D-STED images of samples with STAR-RED, image acquisition was carried out on an Abberior Facility Line STED microscope using a 100x magnification UplanSApo 1.4 NA oil immersion objective lens. Imaging of Abberior STAR RED was done using a pulsed excitation laser of 640 nm and a pulsed 775 nm depletion laser was used. Fluorescence was detected by spectral detectors between 650-763 nm with a time gating delay of 750 ps for an interval of 8 ns. Post-processing of the images was done using the deconvolution software: SVI Huygens Professional (Hilversum, The Netherlands) and further optimization and analyzing of the images were done using FIJI [31].

## Acknowledgements

We thank Jamal-Eddine Bouameur and Thomas Magin at Leipzig University for providing the mice used in this study and Jens Ahm Sørensen, University Hospital Odense, for providing the human epidermis.

We also acknowledge the Danish Molecular Biomedical Imaging Center (DaMBIC, University of Southern Denmark), supported by the Novo Nordisk Foundation (NNF) (grant agreement number NNF18SA0032928), for the use of the bioimaging facilities.

